# *Toxoplasma gondii* infection accelerates the progression of hereditary spastic paraplegia

**DOI:** 10.1101/2024.05.15.594284

**Authors:** James R. Alvin, Carlos J. Ramírez-Flores, Caitlin A. Mendina, Anjon Audhya, Laura J. Knoll, Molly M. Lettman

## Abstract

The parasitic protozoa *Toxoplasma gondii* chronically infects the central nervous system of an estimated one-third of the human population. Infection is generally subclinical, but immunocompromised individuals can experience a variety of neurological symptoms. Meta-analyses of *T. gondii* seropositivity have suggested a correlation between *T. gondii* infection and neurologic disease. While mechanistic studies on the relationship between *T. gondii* infection and neurologic disease have been attempted in mice, mice are particularly susceptible to *T. gondii*, making them an effective model for investigating mechanisms of infection, but not ideal for examining the relationship between long-term chronic *T. gondii* infection and neurologic disease. Rats more closely mimic human clearance of *T. gondii* after acute infection, but a lack of rat models of neurologic disease has limited studies on the interplay between *T. gondii* infection and neurologic disease progression. We have employed a previously characterized rat model of a complex form of hereditary spastic paraplegia (HSP), a class of neurodegenerative disorders which cause axonal degeneration and lower limb spasticity, in order to assess the effect of chronic *T. gondii* infection on neurodegenerative disease. We find that infected rats with hereditary spastic paraplegia exhibit significantly exacerbated behavioral and neuromorphological HSP symptoms compared to uninfected HSP mutant rats, with little correlative effect in infected versus uninfected control animals. We further find that all infected rats regardless of genotype exhibit a robust immune response to *T. gondii* infection, effectively clearing the parasite below the limit of detection of multiple assays of parasitemia and exhibiting no detectable increase in neuroinflammation seven weeks post-infection. These results suggest that chronic undetected *T. gondii* infection may exacerbate neurodegenerative disease even in immunocompetent individuals and may contribute to neurodegenerative disease heterogeneity.

**Author Summary:** The long-term consequences of previous acute infections are poorly understood, but are becoming increasingly appreciated, particularly in the era of long Covid. Altered progression of other diseases later in life may be among the long-term consequences of previous infections. Here we investigate the relationship between previous infection with the parasite *Toxoplasma gondii*, which infects ∼30% of the global population, and neurodegenerative disease using a rat model of hereditary spastic paraplegia (HSP). We find that previous infection with *T. gondii* accelerates motor dysfunction in HSP rats, despite robust clearance of the parasite by infected rats. Our results suggest that previously cleared infections may alter the progression of other diseases later in life and contribute to neurodegenerative disease heterogeneity.

## Introduction

The parasitic protozoa *Toxoplasma gondii* can infect almost any nucleated cell, where it enrobes itself in membrane hijacked from the host cell plasma membrane [1]. Infection with *T. gondii* progresses from an initial acute phase to a chronic phase characterized by the life cycle of the parasite. In the acute phase, fast-growing *T. gondii* tachyzoites disseminate throughout the body, infecting a wide range of tissues. In the chronic phase, starting 2-3 weeks post-infection, most tachyzoites have been cleared by the host cells, but slow-growing bradyzoites, which form intracellular cysts enclosed in an additional carbohydrate shell under the membrane barrier, can persist throughout the life of the host, largely within neurons and muscle [2–4].

Chronic *T. gondii* infection, affecting roughly 30% of the global human population, is generally thought to be asymptomatic in humans with healthy immune systems [5]. However, the persistence of *T. gondii* cysts within axons of chronically infected individuals, reaching tens of microns in size, has raised the possibility of a connection between chronic *T. gondii* infection and neurologic disorders [4,6]. Indeed, some epidemiologic studies have identified a correlation between *T. gondii* infection and the incidence of Alzheimer’s disease [7–10] and neuropsychiatric changes [11–13]. However, other studies have failed to find any correlative relationship between *T. gondii* infection and neurologic disease, and mechanistic information is lacking [14,15].

Due to the heterogeneity in human populations in terms of genetic and environmental modifiers, as well as heterogeneity in terms of strain of *T. gondii* infection and levels of persistent cyst burden, animal studies are needed to unambiguously define mechanistic relationships between *T. gondii* infection and other neurological disorders. Mice are a common model for *T. gondii* studies as they are particularly susceptible to the formation of cysts composed of the slow-growing bradyzoites that persist in muscle and brain tissue during chronic infection [16]. However, this enhanced susceptibility to persistent cysts may thwart their utility for modeling the relationship between chronic *T. gondii* infection and neurologic disease as the high number of intra-neuronal cysts might themselves contribute to long-lasting behavioral changes in mice [11].

Rats are more similar to humans in their response to *T. gondii* infection in that they do not show appreciable sickness during the acute phase and intracellular cysts are sparse during the chronic phase [17–19]. Because their response to *T. gondii* infection is more similar to humans, rats would make a more ideal model to investigate the relationship between long-term chronic infection and neurodegenerative disease, however; a limitation to date has been an absence of rat models that accurately recapitulate human neurodegenerative disease [20,21].

We recently described a rat model of the neurodegenerative disease hereditary spastic paraplegia (HSP) [22]. HSPs constitute a heterogeneous group of neurodegenerative disorders characterized by progressive lower limb spasticity due to functional impairment of the long axons of the corticospinal tract [23,24]. Like many neurodegenerative diseases, genetic causes of HSP are diverse; over 90 genes have been linked to HSP to date [25]. Despite their heterogeneity, HSPs are more genetically tractable than many other neurodegenerative diseases, as in HSP patients whose genomes or exomes have been sequenced, mutations that result in amino acid changes that are hypothesized to contribute to disease can be identified and cross-generational disease incidence follows Mendelian inheritance patterns [26,27].

Patients harboring the p.R106C mutation in TRK-fused gene (TFG), critical for the efficient transport of secreted and membrane-bound proteins from the endoplasmic reticulum to their final destinations, develop severe, early onset HSP [28–30]. We previously used CRISPR/Cas9 genome editing to introduce the p.R106C mutation into the endogenous TFG coding region in Sprague-Dawley rats. Rats homozygous for the p.R106C mutation in TFG recapitulated key aspects of human disease, including progressive gait deficits and corpus callosum deterioration [22]. These HSP rats provided a unique opportunity to investigate the impact of T*. gondii* infection on neurodegenerative disease course in a validated disease model in an organism that resembles the human response to infection.

To determine if *T. gondii* infection accelerates neurodegeneration, we infected wild-type Sprague-Dawley rats and rats homozygous for the p.R106C mutation in TFG (hereafter termed HSP rats) and compared their gait, behavior, brain structure, and transcriptional profiles to their uninfected counterparts. We show that despite persistent seropositivity to *T gondii* antigens, cysts are rare or absent in wild-type or HSP mutant brains. Additionally, despite the absence of a continued high burden of *T. gondii*, infected HSP rats exhibit accelerated motor and neuromorphological phenotypes relative to their uninfected counterparts. These behavioral and structural changes were not due to persistent neuroinflammation in response to HSP or infection. These results suggest that *T. gondii* can exacerbate and accelerate neurodegeneration and highlight the potential contribution of infectious agents with brain tropism to neurological disorders.

## Materials and Methods

All animal studies were approved by the Institutional Animal Care and Use Committee of the University of Wisconsin-Madison. HSP rats were previously described and carry two copies of the p.R106C mutation in TFG [22].

### *T. gondii* oral infection

Eight Swiss Webster mice were injected with 1×10^4^ tachyzoites. After 4-5 weeks, their brains were harvested and fed to six-week-old wild-type and HSP Sprague-Dawley rats (four each). Each rat received approximately 5,000 cysts. Brains from uninfected mice were fed as a mock infection to two HSP rats.

### *T. gondii* infection via intraperitoneal injection

Five wild-type and five HSP mutant Sprague-Dawley rats underwent intraperitoneal injection of 2.5×10^8^ tachyzoites at six weeks of age. Three wild-type and three mutant Sprague-Dawley rats were mock injected with saline.

### Quantitative gait measurements

Gait was assessed as described previously [22]. Briefly, animals were recorded as they traversed a clear platform and positional coordinates of points of interest were assigned by a pre-trained neural network. Kinematic parameters were calculated using custom MATLAB scripts.

### Quantitative behavior assessment (Open Field)

Animals were placed under camera observation in an open box for 30 minutes undisturbed. The 30-minute period was divided into three 10-minute intervals, over which animal movement was analyzed with ANY-maze software.

### Analysis of brain sections

Animals were anesthetized with isoflurane and transcardially perfused with PBS followed by 4% paraformaldehyde (PFA). Five-micron coronal sections of paraffin-embedded tissue were deparaffinized and rehydrated. For histological analysis, tissues were stained with 0.1% Luxol fast blue, hematoxylin, and eosin. Images were acquired using a uScope HXII slide scanner and regions of interest were measured with ImageJ. For immunofluorescence and quantification of astrocyte and microglia density, antigen retrieval was performed on deparaffinized sections. After blocking, tissues were incubated with primary antibodies overnight (Iba1 (019-19741, FUJIFILM Wako Pure Chemical Corporation) or S100b (GA50461-2, Agilent)), washed and incubated with secondary antibodies and mounted with DAPI. Images were acquired with a Nikon Eclipse Ti2-E spinning disk confocal microscope equipped with a Yokogawa CSU-W1 scanhead and an ORCA-Fusion BT sCMOS camera at 20x magnification. Positively stained cells in a 1500 x 1500 pixel square in the primary motor region were counted with Imaris (Oxford Instruments).

### Cyst quantification

Ground brain samples were stained with biotinylated Dolichos biflorus agglutinin (B-1035-5; Vector Laboratories) and serum from a mouse with a chronic *T. gondii* infection, washed, and incubated with Strepavidin-conjugated Alexa Fluor 488 and anti-mouse Alexa Fluor 594 (Thermo Fisher). An aliquot of processed brains was mounted on a glass coverslip and cysts were visualized with a Zeiss Axioplan III motorized microscope with a 40× objective.

### In vivo imaging system (IVIS)

Seven Sprague-Dawley rats heterozygous for the TFG p.R106C mutation were infected via intraperitoneal injection at nine weeks of age with a strain of *T. gondii* engineered to express firefly luciferase [31]. Five rats received 10^7^ *T. gondii* tachyzoites and two received 10^8^ tachyzoites. Animals were euthanized by CO_2_ asphyxiation at 13 weeks and their brains were placed in cold PBS. Brains were soaked in 10 mg/mL D-Luciferin potassium salt (15.4 mg/mL in PBS) for five minutes prior to imaging with an *in vivo* imaging system (IVIS, PerkinElmer).

### Genomic DNA amplification

Animals were euthanized by CO_2_ asphyxiation. Brains were harvested and the cortex was dissected on ice and flash frozen in TRIzol (Invitrogen). Tissue was subjected to bead homogenization in TELT lysis buffer (50 mM Tris HCl, 62.5 mM EDTA, 4% Triton X-100, 2.5 M LiCl) and genomic DNA was purified using phenol-chloroform extraction. Control DNA was prepared from tissue culture-derived tachyzoites. Genomic DNA was amplified with GoTaq (Promega) using *T. gondii* or *R. norvegicus* specific primers.

*T. gondii* SAG1: TGCCCAGCGGGTACTACAAG, TGCCGTGTCGAGACTAGCAG

*T. gondii* Tubulin-M: CCAACCTGAACAGACTGATTGCC, TTGGTCTGGAACTCAGTCACGTC

*T. gondii* GAPDH: ATGCTTAACGACACCTTCGTTAAGC, CCTGGACGGACATGTAGTGAG

*R. norvegicus* GAPDH: AACCCATCACCATCTTCCAG, CCAGTAGACTCCACGACATAC

### Western blot

To monitor the presence of *T. gondii* antibodies in serum of infected animals, whole tachyzoites were lysed in RIPA buffer (50 mM Tris HCl, 150 mM NaCl, 1.0% (v/v) NP-40, 0.5% (w/v) Sodium Deoxycholate, 1.0 mM EDTA, and 0.1% (w/v) SDS a pH of 7.4). One hundred micrograms of protein were separated on a 10% acrylamide gel and transferred to a nitrocellulose membrane. The membrane was cut in strips and blocked in PBS-Tween 5% milk, then incubated with serum from pre-infected, orally infected, and intraperitoneally infected wild-type and HSP mutant rats (1:1000 in PBS-1% Tween) followed by secondary antibody (1:10,000 goat anti-rat HRP), and revealed with ECL prime Western Blotting detection reagent (Cytiva). The membrane was imaged on a Licor imager.

### RNA isolation and sequencing

Infected and uninfected wild-type and HSP mutant animals were euthanized by CO_2_ asphyxiation. Brains were harvested and the region containing the motor cortex was dissected on ice and flash frozen in TRIzol. RNA was extracted using phenol/chloroform separation and isopropanol precipitation. RNA samples were DNase treated. The purity of nucleic acids was confirmed by A260/A280 and A260/A230 ratio range 1.8 - 2.1 and one single peak directly over 260nm.

The University of Wisconsin-Madison Biotechnology Center’s Gene Expression Center Core Facility (research resource identifier [RRID]: SCR_017757) conducted quality control assessments, RNA library preparation, sequencing, and read demultiplexing. Library construction involved poly(A) enrichment using the Illumina TruSeq stranded mRNA kit. Double stranded cDNA was purified by AMPure Xp beads and frozen at −20C. Library quantification was performed on Agilent Biomek Synergy H1 Plate Reader with Picogreen reagent and final libraries assayed on Agilent 4200 Tapestation with HS D1000 ScreenTapes. Following quality control, samples were sequenced on the NovaSeq S4 platform (Illumina) with a 2 × 150-bp configuration. On average, each sample generated approximately 50 million paired-end reads, with an average read length of 450 base pairs. Raw RNA sequencing data have been deposited in NCBI and are accessible through the BioProject identification number: PRJNA1110596.

### Transcriptome assembly

RNA sequencing analysis was performed as previously reported with the following changes [32]. Briefly, the sequencing reads were processed to remove low quality reads using Trimmomatic, v0.39 [33]. Reads for all samples were aligned to the Rattus norvegicus genome assembly mRatBN7.2 (https://useast.ensembl.org/ and https://www.ncbi.nlm.nih.gov/, respectively) using the Spliced Transcripts Alignment to a Reference program (STAR, v2.7.0f) [34]). The default STAR parameters were selected excepting the following settings: (a) mismatch of 2 bp and (b) intron length (20-100,000 bp). Quantification of mapped reads and the generation of a counts table were conducted (RSEM v1.3.1) [35]. Counts were imported into R (tximport v1.16) [36] and differential expression analysis was conducted using DESEq2 v1.28.1 [37]. The log-transformed DESeq2 values were used for the generation of PCA plots in the DESeq2 package. Only genes whose differential expression met a *p*-value cutoff of <0.05 in a t test and with IDs assigned in the Rat Genome Database were included in subsequent analysis.

### Gene Ontology classification

Genes whose expression changed by at least 30% upon infection in wild-type animals were entered into DAVID functional annotation tools to classify genes based on their biological process. Upregulated and downregulated lists were entered separately. Groups containing at least 7 differentially expressed genes were plotted. DAVID, Database for Annotation, Visualization and Integrated Discovery [38,39].

## Results

### *T. gondii* infection accelerates motor decline in HSP animals

While recent epidemiological work has linked toxoplasmosis to neurodegenerative disease [7,8,10], the mechanistic relationship between *T. gondii* infection and neurodegenerative disease progression has not been examined outside of mice, which are much more susceptible to *T. gondii* infection than humans [16]. We have previously shown that Sprague Dawley rats homozygous for the HSP-associated p.R106C mutation in TFG recapitulate key aspects of human disease progression, including progressive motor decline and a thinning corpus callosum [22]. We leveraged this model to assess the impact of *T. gondii* infection on neurodegenerative disease progression.

To first establish *T. gondii* infection conditions and evaluate acute and chronic responses to infection in Sprague Dawley rats, we orally infected animals at 6 weeks of age with 5,000-10,000 cysts. These animals exhibited no symptoms during the initial acute infection and some animals fed cysts failed to exhibit detectable *T. gondii* antibodies in their serum at approximately 13 weeks of age (Fig. S1A). We therefore infected a second cohort of animals via intraperitoneal injection with luminescent *T. gondii* and assessed infection postmortem (Fig. S1B) [31,40]. Despite only one brain from an injection infected animal displaying luciferase activity indicating the presence of *T. gondii*, and too few cysts to perform accurate whole-brain quantification and no detectable *T. gondii* genomic DNA in their brain tissue (Fig. S1B, S1C, S1D), animals infected via injection displayed a more robust presence of *T. gondii* antibodies in their serum at approximately 13 weeks of age (Fig. S1A). As humans with chronic *T. gondii* infection typically maintain *T. gondii* antibodies in their serum with little/no detectable evidence of persistent cysts in their brains, our infected rats recapitulated human infection [41]. Because it was not readily apparent if one infection route would more substantially contribute to behavioral dysfunction, animals infected via both routes were subject to further analysis; only animals lacking detectable *T. gondii* antibodies in their serum were excluded (Fig. S1A).

We previously showed that HSP mutant animals have quantifiable gait deficits at 13 weeks of age [22]; therefore, we assessed gait function in our chronically infected animals at 13 weeks of age to determine if *T. gondii* infection exacerbates motor dysfunction (Fig. 1A, 1B). Consistent with our previous data [22], 13-week uninfected HSP mutant animals exhibited exaggerated lateral oscillation of the hindbody, as measured by hindbody sway (Figure 1B, 1C). *T. gondii* infection significantly exacerbated lateral hindbody oscillation of HSP mutant animals with no effect on their wild-type counterparts (Fig. 1C). HSP mutant rats are also unable to maintain normal tail height (Fig. 1B, 1D) [22]. *T. gondii* infection did not impact tail height in HSP mutant rats; even uninfected mutant animals hold their tails very close to the ground with little room for further dropping of the tail (Movie S1). Interestingly, *T. gondii* infection decreased the ability of wild-type rats to maintain normal tail height (Figure 1D). Overall, we find that *T. gondii* infection significantly exacerbates motor decline in HSP mutant rats, as well as impacting normal motor function in wild-type animals.

**Figure. 1.**
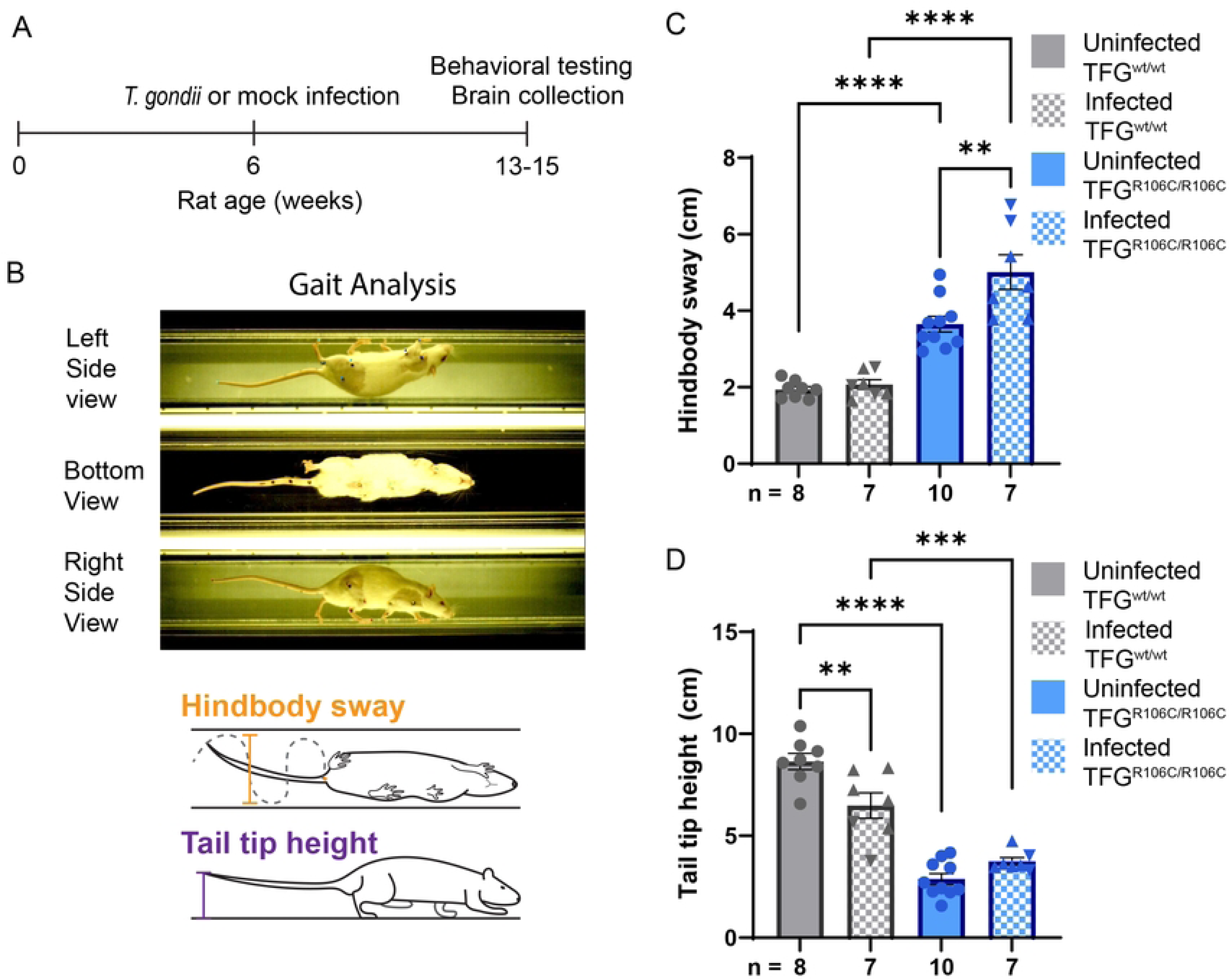
*T. gondii* infection accelerates motor decline in HSP rats. (A) Timeline of infection, and behavioral and brain assessments. (B) Single frame of video used for gait analysis. Colored points indicate software-assigned positions (*Top*). Schematics illustrating hindbody sway and tail tip height quantification (modified from [22]) (*Bottom*). (C-D) Measurements of hindbody sway (C) and tail tip height (D) of animals of the indicated genotype and infection condition. Data points represent average measurements from individual animals. Triangle symbols represent data points from animals infected via injection, while inverted triangle symbols represent data from animals infected orally. Error bars represent mean ± SEM. **P < 0.01, ***P < 0.001, and ****P < 0.0001, as calculated using Tukey’s multiple comparisons test. wt, wild-type; ns, non-significant.

Previous assessments of the impact of *T. gondii* infection on behavior in rodents have suggested that infected animals are more active and exhibit fewer responses associated with fear [42,43]. To investigate the impact of *T. gondii* infection on overall activity levels, we observed infected and uninfected wild-type and HSP animals in a 30-minute open field test (Fig 2A). As overall activity decreases over time as animals adapt to their new environment, the 30-minute test was divided into three 10-minute periods [44]. We assessed the mean speed and distance traveled as measures of overall activity, as well as time spent freezing, defined in our experiments as the lack of any frame-to-frame pixel differences, as an additional measure of activity and a potential correlate of fear activity. Interestingly, infected wild type animals exhibited no significant difference from their uninfected counterparts in mean speed, distance traveled, or time spent freezing in any of the three time periods (Fig. 2B, C, D). Although no changes reached statistical significance, HSP mutant rats exhibited an overall trend toward reduced activity with reduced mean speed and distance traveled, and increased time spent freezing in all three time periods (Fig. 2B, C, D). This is consistent with our qualitative observations that infected HSP mutant animals moved less in their cages and upon handling. Interestingly, the only pairwise comparisons that reached statistical significance were between wild-type and HSP mutant infected animals in some of the time points in all three measures of activity (Fig. 2B, C, D). This suggests that, although we were unable to measure a difference in activity between infected and uninfected wild-type animals, infection may result in a modest increase in activity in the absence of pre-existing neurodegenerative insults, consistent with some previous reports [45].

**Fig. 2:**
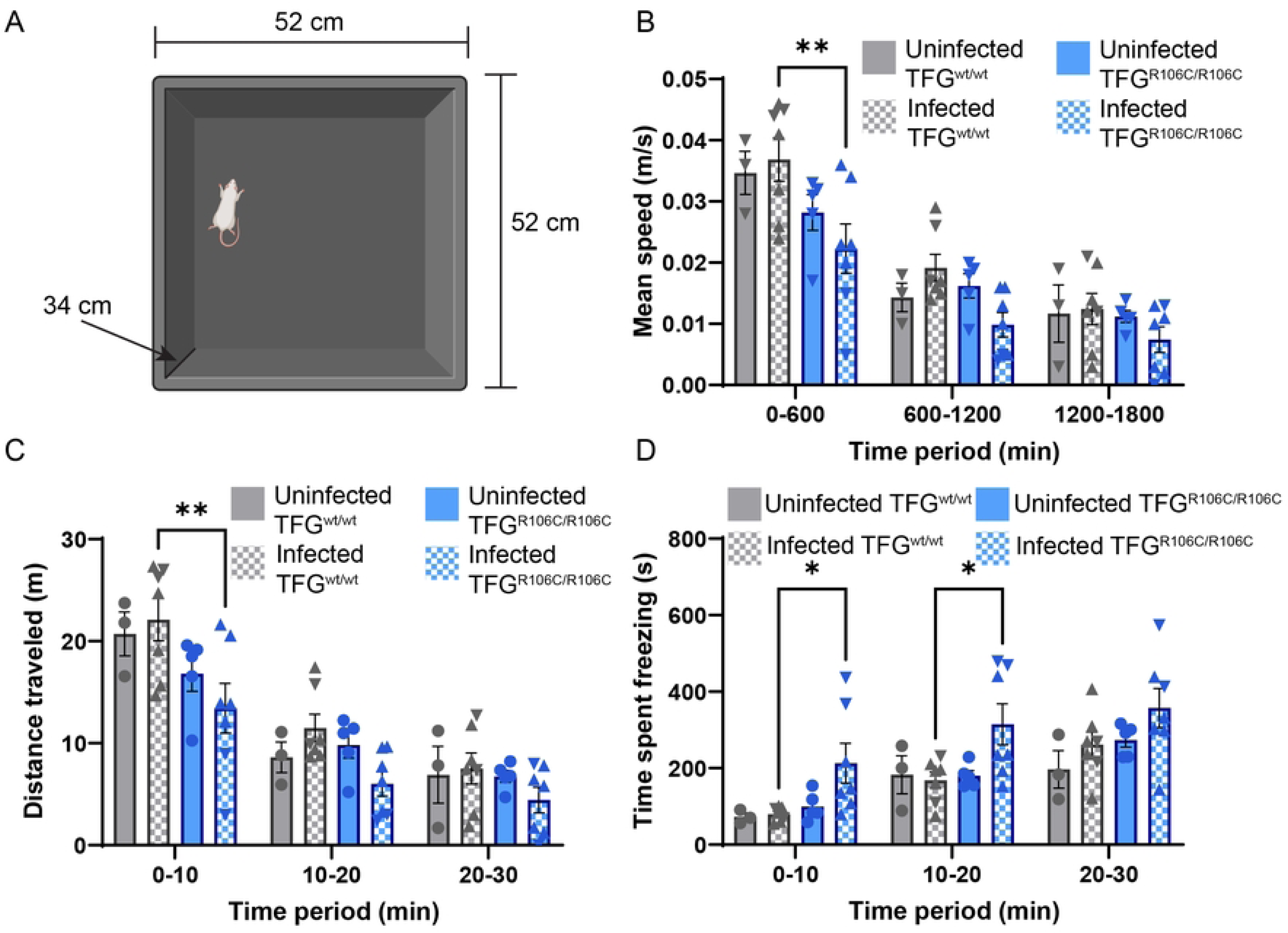
*T. gondii* infection alters behavior in HSP mutant animals. (A) Schematic illustrating the open field test (created with Biorender.com). (B-D) Mean speed (B), total distance traveled (C), and time spent freezing in place with no frame-to-frame pixel differences (D) in the open field test of animals of the indicated genotype and infection condition. Each data point represents an individual animal. Uninfected wild-type, n = 3; infected wild-type, n = 7; uninfected HSP mutant, n = 5; infected HSP mutant, n = 7. Triangle symbols represent data points from animals infected via injection, while inverted triangle symbols represent data from animals infected orally. Error bars represent mean ± SEM. *P < 0.05 and **P < 0.01, as calculated using Tukey’s multiple comparisons test. wt, wild-type.

### *T. gondii* infection accelerates CNS pathology in HSP animals

Despite heterogeneity in terms of underlying mutation, MRI studies have identified thinning of the corpus callosum and ventriculomegaly as common features of HSP patients [46,47]. Our rodent model exhibits progressive corpus callosum thinning consistent with human findings [28–30]. Although they are indistinguishable at 13 weeks of age, by 25 weeks, the corpus callosum of HSP mutant animals is significantly smaller than that of wild-type animals [22]. To assess the impact of infection on brain morphology, we stained paraffin-embedded coronal sections using Luxol fast blue (LFB) to mark myelinated axons in the corpus callosum and hematoxylin and eosin (H&E) to allow easy visualization of other brain regions. By tracing the outline of the corpus callosum in coronal brain sections (Fig. 3A), we confirmed that HSP mutant animals have a corpus callosum size similar to wild-type animals at 13 weeks of age, consistent with our previous results (Fig. 3B) [22]. *T. gondii* infection did not alter corpus callosum area in wild-type animals (Fig. 3B). However, in *T. gondii* infected HSP mutant animals, corpus callosum area was significantly reduced compared to both infected wild-type and uninfected HSP mutant animals (Fig. 3B). Despite a trend toward larger ventricles in HSP mutant animals, no significant difference between ventricle size was identified between groups, including no difference due to infection (Fig. S2A).

**Fig. 3.**
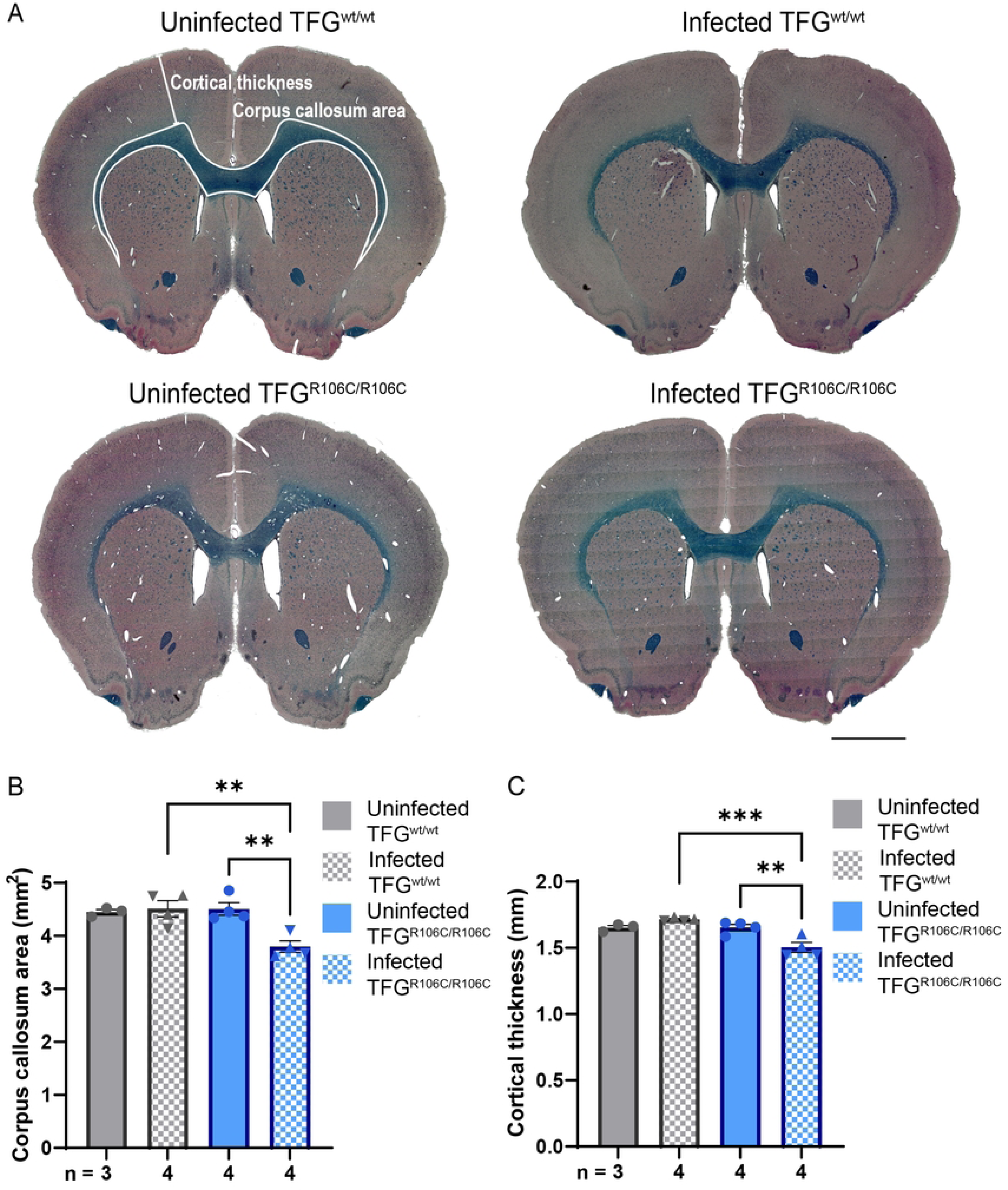
*T. gondii* infection accelerates CNS pathology in HSP rats. (A) Representative images of paraffin sections from 13-week-old animals stained with LFB and H&E from animals of the indicated genotype and infection condition. Representative measurements of cortical thickness and corpus callosum area are marked (*Top Left*). Scale bars, 2mm. (B-C) Quantification of cortical thickness (B) and corpus callosum area (C) of animals of the indicated genotype and infection condition. Data points represent measurements from individual animals. Triangle symbols represent data points from animals infected via injection, inverted triangle symbols represent data from animals infected orally. Error bars represent mean ± SEM. *P < 0.05, **P < 0.01, and ***P < 0.001, as calculated using Tukey’s multiple comparisons test. wt, wild-type; ns, non-significant; LFB, luxol fast blue; H&E, hematoxylin and eosin.

Because *T. gondii* infected HSP rats exhibited increased motor dysfunction relative to uninfected HSP animals, we examined the primary motor cortex where the cell bodies of the upper motor neurons reside. We measured the shortest linear distance between the corpus callosum and the dorsal edge of the cortex (Figure 3A). As with corpus callosum area, infected wild-type and uninfected wild-type and HSP mutant animals did not significantly differ in this parameter (Fig. 3C). In contrast, infected HSP mutant rats exhibited a modest but significantly reduced cortical thickness compared to both infected wild-type and uninfected mutant animals (Fig. 3C).

Because animals infected via intraperitoneal injection exhibited increased prevalence of *T. gondii* antibodies in their serum, we questioned whether animals infected via different routes exhibited differences in their behavioral or CNS pathology. Despite the higher presence of *T. gondii* antibodies in serum of animals infected intraperitoneally (Fig. S1A), both wild-type and HSP mutant animals infected orally exhibited more pronounced differences in every behavioral measure that was different between groups (Fig. S2 B-F). Different infection routes did not result in a difference in CNS pathology, however (Fig. S3 A,B).

### Chronic *T. gondii* infection does not result neuroinflammation in rats

Various forms of neurodegeneration have been linked to chronic neuroinflammation and chronic *T. gondii* infection leads to a robust and long-lasting immune response in brains of infected mice; we therefore hypothesized that neuroinflammation in chronically infected HSP rats may exacerbate motor dysfunction [48–51]. To assess whether *T. gondii* infection exacerbated the neuroinflammatory response in HSP mutant rats, we quantified astrocytes and microglia in the motor cortex of coronal brain sections. We previously showed that the density of both astrocytes and microglia increase in 25-week-old, but not 13-week-old, HSP rats [22]. Consistent with our previous results, we found no significant increase in astrocyte or microglia density in uninfected HSP rats relative to wild-type animals (Fig. S3C,D). Surprisingly, infection did not result in an increase in astrocyte or microglia density in wild-type or HSP mutant animals (Fig. S3C,D).

Although we failed to detect signs of neuroinflammation in our cell counting experiments, we wondered if a more sensitive assay would reveal low levels of neuroinflammation or other perturbations that contribute to motor dysfunction. Because the cell bodies of the upper motor neurons implicated in HSP reside in the primary motor cortex, and we hypothesized that changes in this region would most strongly contribute to motor dysfunction observed in uninfected or infected HSP animals, we isolated RNA from this region for RNA sequencing analysis. All extracted RNA was of sufficiently high quality for sequencing. However, upon principal component analysis of the sequencing data, we found that one of our infected mutant samples substantially differed from the others, leading us to exclude it from further analysis (Sample 9, Fig. S4).

We initially focused our attention on genes whose expression changed greater than 2-fold in infected relative to uninfected rats (expression relative to uninfected <0.50 or greater than 2.0). In wild-type *T. gondii* infected rats, 21 genes met this cutoff, while in HSP mutant rats, only six genes were differentially expressed upon infection (Fig. 4A). In striking contrast to robust inflammatory transcriptional changes present up to at least 180 days post infection in brains of *T. gondii* infected mice, both *T. gondii* infected wild-type and HSP rats lacked transcriptional changes suggesting robust, prolonged inflammation (Fig. 4B) [49]. When we examined all genes that were differentially expressed upon *T. gondii* infection in wild-type animals, not restricting ourselves to those that were changed greater than 2-fold, we again did not see strong inflammatory signatures (Fig. S5, Table S1), despite the prolonged seropositivity to *T. gondii* antibodies we observed (Fig S1A).

**Fig. 4:**
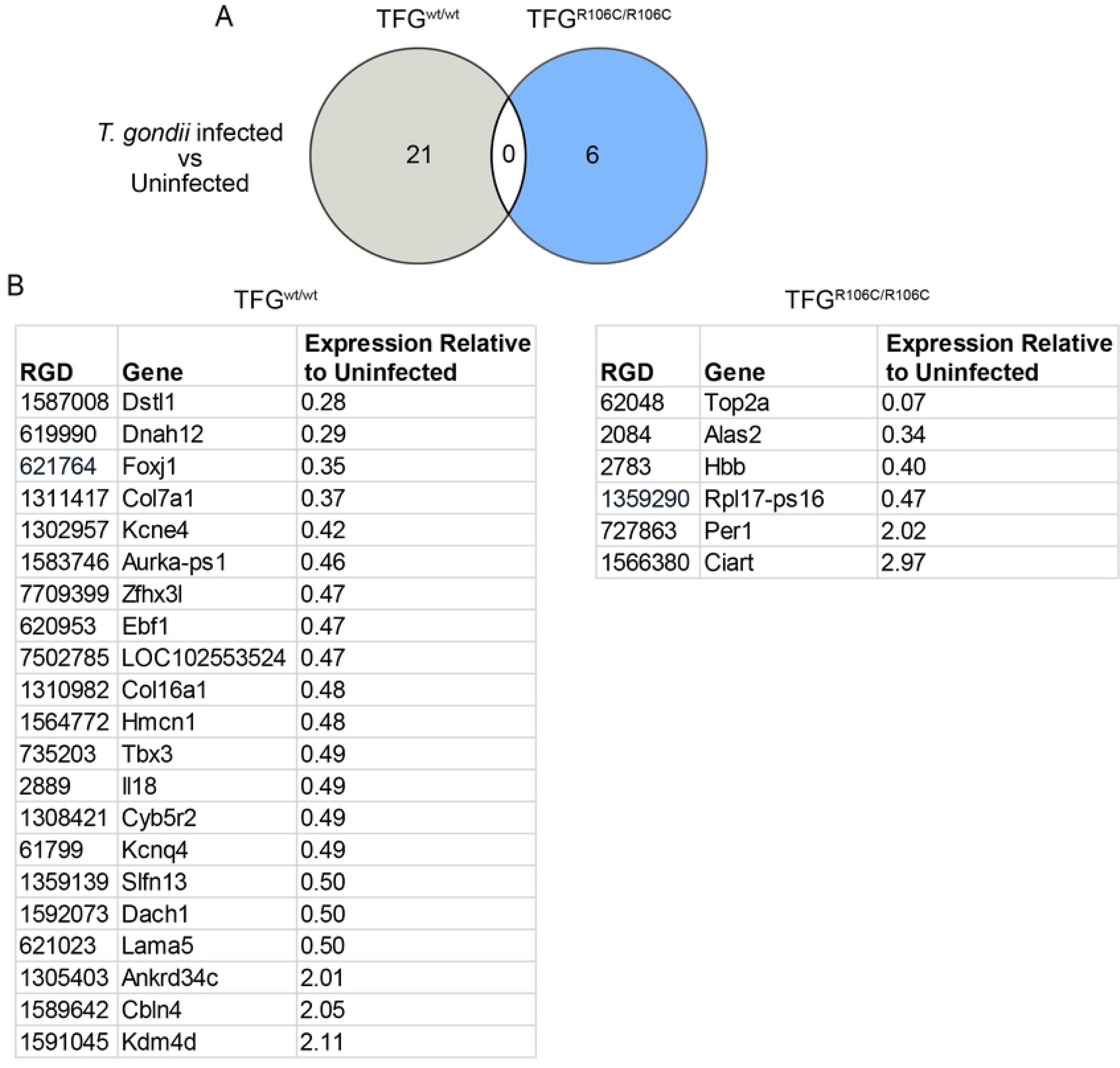
RNA sequencing of HSP mutant and wild-type animals reveals little evidence of persistent inflammation following *T. gondii* infection. (A) Venn diagram comparing the genes differentially expressed in TFG wild-type and mutant brains with differential expression defined as a greater than 2-fold change (expression relative to uninfected <0.50 or greater than 2.0). (B) List of genes differentially expressed upon *T. gondii* infection in TFG wild-type (*Left*) or TFG mutant (*Right*). wt, wild-type; RGD, Rat Genome Database.

Surprisingly, there was no overlap between the genes differentially expressed upon *T. gondii* infection in wild-type and HSP mutant brains (Fig. 4B) and similarities were not readily apparent. The small number of genes differentially expressed upon infection in HSP mutant animals precluded robust gene ontology analysis (Table S1).

Interestingly, the three genes most highly upregulated upon infection in wild-type brains (Ankrd34c, Cbln4, Kdm4d), as well as several downregulated genes (Col7a1, Col16a1, and Lama5) may influence remodeling of or are themselves extracellular matrix proteins, suggesting that long-lasting effects of *T. gondii* infection, even in the absence of persistent cysts, may arise either through continued remodeling of the ECM to recover and rebuild after infection, or due to continued circulating *T. gondii* antibodies [52–54].

The only two genes upregulated more than 2-fold upon infection in HSP mutant brains are both involved in regulating circadian rhythms (Per1 and Ciart). Interestingly, Per1 has been shown to regulate the immune response to prevent excessive macrophage recruitment, raising the possibility that the upregulation of these genes protects animals from rampant neuroinflammation [55,56].

Consistent with our inability to identify evidence for persistent *T. gondii* cysts by histological examination or PCR of genomic DNA (Figs. 3A, S1D) we did not detect *T. gondii* genes in our RNA sequencing analysis. This is consistent with either undetectable or low levels of *T. gondii* DNA and RNA in postmortem human samples despite global seropositive rates of ∼25-30% [5,15,57–59].

## Discussion

With approximately 30% of the global population harboring latent *T. gondii*, and increasing rates of neurodegenerative disease, often with unknown etiology, we hypothesized that there may be a relationship between *T. gondii* infection and neurodegenerative disease [5,60]. Indeed, previous epidemiologic studies have suggested a correlation between *T. gondii* infection and neurodegenerative disease [7–10]. While mechanistic explorations of this relationship have been attempted in mice, the increased propensity for *T. gondii* cysts to persist in mouse brains relative to humans have made it difficult to extend the results of these studies to human disease [16].

Here we utilized a validated rat model of the motor neuron disease hereditary spastic paraplegia [22]. As, similar to humans, *T. gondii* cysts are rare in rat brains after acute infection [17–19], this model allowed us to probe the impact of chronic *T. gondii* infection on neurodegenerative disease without the confounding presence of a high number of cysts. Indeed, our infected rats exhibited no acute symptoms and effectively cleared *T. gondii* from the brain. Despite low or undetectable *T. gondii* cysts in the brains of previously infected animals, *T. gondii* antibodies persisted in serum, an outcome consistent with humans in the chronic phase of infection [41]. Strikingly, we found that prior *T. gondii* infection modestly, but significantly exacerbated motor dysfunction in HSP rats, as measured by an increase in side to side hindbody movement and reduced overall locomotor activity of infected HSP rats. Chronically infected HSP rats had a modest, but significant decrease in thickness in the primary motor area and a decrease in corpus callosum thickness, neuromorphological changes that potentially underlie the exacerbated motor dysfunction.

When we sought to understand the molecular changes underpinning the exacerbated motor dysfunction in HSP animals by transcriptional analysis, we found that the expression of relatively few genes was altered in infected HSP animals, leaving few clues into the mechanism underlying the relationship between chronic *T. gondii* infection and progression of neurodegenerative disease. The upregulation of genes involved in circadian rhythms is striking in the absence of a large number of other transcriptional changes in response to infection. Changes in the circadian system have been previously correlated with neurodegenerative disease [61–63] and with regulating the immune response [55,56,64,65], suggesting that these pathways, or the interplay between them, may contribute to accelerated motor dysfunction in *T. gondii* infected HSP rats.

Contrary to our initial hypothesis that *T. gondii* and HSP may interact via neuroinflammation, we detected no evidence of a heightened immune response by either inflammatory cell counts (Fig. S3C, S3D) or transcriptional analysis (Fig. 4). While chronically infected HSP rats do not exhibit persistent neuroinflammation, they do maintain circulating *T. gondii* antibodies. One possibility is that continued circulating cytokines impact accelerated motor dysfunction in infected HSP mutant animals [66,67]. Indeed, previous exposure to viral pathogens, including viruses without known brain tropism, has been linked to increased risk of neurodegeneration in humans [68]. Alternatively, potential damage to or loss of neurons during the clearance of the acute infection may be responsible for accelerated motor dysfunction.

In both infected groups, we were surprised by the lack of transcriptional changes related to an inflammatory response. Previous work in mice has demonstrated robust and prolonged inflammation in brains of infected mice [48,49]. This absence of prolonged neuroinflammation may be explained by the robust clearance of *T. gondii* by rats, as opposed to mice. We attempted five methods of cyst detection: examination of brain sections (Fig. 3A), imaging of cyst bioluminescence in brains of animals infected with *T. gondii* expressing luciferase (Fig. S1B), counting cysts in dissociated brain tissue (Fig. S1C), RNA sequencing (Fig. 4), and genomic DNA quantification (Fig. S1D). None of these methods returned more than trace evidence of *T. gondii* cysts in infected brains, further confirming that rats, similar to humans [17–19], largely clear *T. gondii* from their brains.

Previous experiments have suggested a prolonged impact of chronic *T. gondii* infection on animal behavior, even in the absence of underlying mutations linked to neurodegenerative disease. This is suggested by epidemiologic studies correlating chronic *T. gondii* infection and neuropsychiatric disorders, as well as more common place behaviors such as risk-taking [11–13,58]. In open field tests, *T. gondii*-infected mice typically demonstrate hyperactive behavior expressed through increased distance traveled and average speed [45]. Chronically infected wild type rats did exhibit a modest, but significant, change in motor function. Infected wild type rats did not maintain tail height consistent with uninfected animals (Fig. 1B) and, while not significantly different, there was a trend toward increased locomotor activity in infected wild type animals (Fig. 2B, C). Increased locomotor activity in rodents and reduced fear response, as suggested by previous studies examining rodent response to predator scents, have been suggested to make infected rodents more vulnerable to predation and enable the spread of *T. gondii* through ingestion of brain cysts.

Studies examining host response to *T. gondii* infection vary in the reported host outcomes, even within the same host species. This may result from differences in infection modality, infectious dose, *T. gondii* strain, and how *T. gondii* was propagated prior to infection. Here we infected animals orally with ∼5000 tissue cysts or intraperitoneally with 250 million tissue culture-derived tachyzoites. Even at these extremely high doses, tissue cysts were largely undetectable. Our initial assessment of *T. gondii* infection by serum antibody detection led us to consider two wild-type and two HSP mutant animals which underwent oral infection treatment to potentially not have been infected with *T. gondii* to the same extent as other animals and therefore discounted from the study (Fig. S1). We then employed intraperitoneal injection to infect subsequent cohorts of animals. Though *T. gondii* antibodies in serum of animals infected via intraperitoneal injection was generally higher than in animals that ingested bradyzoite cysts (Fig. S1), we found that in all of our behavioral metrics, animals infected orally exhibited more severe symptoms than animals infected via injection (Fig. S2), though there was no corresponding significant difference in brain architecture or inflammatory cell counts (Fig. S3A-D). While this difference between oral and intraperitoneal infection may be interesting to explore in the future, it could also be that we imposed a selection bias for more susceptible animals by removing the four animals with a lower antibody titer from the dataset, leading to the observed behavioral differences between oral and intraperitoneal infection.

The absence of a strong immune response in the transcriptional analysis of wild type chronically infected rats allowed us to observe other changes that may function in parasite clearance or remodeling after infection and may impact long term behavioral changes in response to infection. In particular, the transcription of several genes involved in the function or regulation of the extracellular matrix were altered in chronically infected wild type animals. The role of the extracellular matrix in long-lasting response to prior infection, beyond *T. gondii*, warrants further study [69,70].

Together our results justify increased awareness and investigation of the interplay between infectious and neurodegenerative disease. Even in our rats with undetectable long-term persistence of *T. gondii* in the brain, an acceleration of neurodegenerative disease was measurable, suggesting that differences in neurodegenerative disease severity in human populations may be due, at least in part, to differences in infectious disease burden, and public health initiatives aimed to improve hygiene for underserved populations may improve long term health outcomes for more reasons that previously appreciated.

## Acknowledgments

Support was also provided by the UWCCC Genome Editing and Animal Models Shared Resource, the UWCCC Flow Cytometry Laboratory, and the UW Optical Imaging Core facility. We thank members of the Audhya and Knoll labs for useful discussions and the Waisman Center Rodent Models Core staff for their excellent animal care.

## Author contributions

Conceptualization: J.R.A., C.J.R.F., A.A., L.J.K., and M.M.L.

Investigation: J.R.A., C.J.R.F., C.A.M., and M.M.L.

Formal analysis: J.R.A., C.J.R.F., and M.M.L.

Writing-original draft: J.R.A.

Writing-review & editing: J.R.A, C.J.R.F., C.A.M., A.A., L.J.K., and M.M.L.

Funding Acquisition: A.A. and L.J.K.

---

**Figure S1. Infected animals exhibit minimal brain cyst presence.** (A) Western blot strips from *T. gondii* antibody titer test. *T. gondii* proteins were separated via SDS-PAGE, transferred to nitrocellulose, incubated with the indicated rat sera, and exposed to anti-rat HRP. (-) = secondary antibody only. Strips marked with * denote animals deemed incompletely infected and excluded from further analysis. (B) Representative *in vivo* imaging system (IVIS) results of infected brains of heterozygous TFG p.R106C Sprague-Dawley rats following treatment with 10 mg/mL luciferin. The five brains on the upper two rows were harvested from animals injected with 10^7^ tachyzoites, while the two brains on the lower row were from animals injected with 10^8^ tachyzoites. Radiance scale on left. (C) Representative images of a *T. gondii* cyst harvested from the circled brain in Fig. S1B. Brain tissue was homogenized and stained using serum from a mouse with a chronic *T. gondii* infection (*top left*, red) and Dolichos biflorus agglutinin (*top right*, green). Brightfield images of the cyst and encased parasites (*bottom left*, gray) and merge (*bottom right*). Scale bar, 20 μm. (D) PCR of gDNA. Lanes 3, 5, 7, 12, and 14-16 are wild-type rat gDNA; lanes 4, 8-10, 13, and 17-19 are HSP mutant rat gDNA. bp, base pairs

**Fig. S2. Orally infected animals exhibit more severe symptoms than animals infected via intraperitoneal injection.** (A) Ventricle area was not altered in infected animals. (B,C) Gait measurements of hindbody sway (B) and tail tip height (C) of animals of the indicated genotype and infection method. Data points represent measurements from individual animals. Error bars represent mean ± SEM. *P < 0.05 and **P < 0.01, as calculated using Tukey’s multiple comparisons test. wt, wild-type. (D-F) Measurements of distance traveled (D), average speed (E), and time spent freezing (F) in the open field test by animals of the indicated genotype and infection condition. Data points represent measurements from individual animals. Orally infected wild-type, n = 2; intraperitoneally (IP) infected wild-type, n = 5; orally infected HSP mutant, n = 2; IP infected HSP mutant, n = 5. Error bars represent mean ± SEM. *P < 0.05 and **P < 0.01, as calculated using Tukey’s multiple comparisons test. wt, wild-type.

**Fig. S3. Brain morphology did not differ between infection conditions; *T. gondii* infection did not cause gliosis.** (A-B) Measurements of corpus callosum area (A) and cortical thickness (B) of animals of the indicated genotype and infection method. Data points represent measurements from individual animals. Error bars represent mean ± SEM. (C-D) Measurements of astrocyte (C) and microglia (D) density in brains of animals of the indicated genotype and infection condition. Data points represent measurements from individual animals. Error bars represent mean ± SEM.

**Fig. S4. An outlier infected TFG mutant sample was excluded from RNA sequencing analysis.** PCA plot of RNA sequencing data. Brain tissue from the motor cortex was snap frozen in TRIzol and RNA was isolated. High-quality RNA was submitted for RNA sequencing. Based on PCA analysis, sample 9 was excluded from further analysis. PCA, principal component analysis.

**Fig. S5. Gene ontology classification of genes differentially regulated upon *T. gondii* infection in wild-type animals.** DAVID gene ontology tools were used to classify genes differentially expressed in the primary motor region of wild type animals. Genes whose expression differed at least 30% upon infection were included in the analysis. Groups containing at least 7 differentially expressed genes are shown. wt, wild-type; DAVID, Database for Annotation, Visualization and Integrated Discovery.

**Table S1. List of genes differentially expressed in the primary motor region of wild-type animals upon chronic *T. gondii* infection.** Genes whose expression differed at least 30% upon infection are shown. RGD, Rat Genome Database.

**Movie S1. Representative videos of 13-week male rats crossing a transparent platform.** A camera underneath the platform uses mirrors to capture three views of rat motion. Colored points added by analysis software. Uninfected TFG wild-type (*top left*), uninfected TFG mutant (*top right*), infected TFG wild-type (*bottom left*), and infected TFG mutant (*bottom right*) animals are shown. Playback speed is 0.5x real time.

